# SnFFPE-Seq: towards scalable single nucleus RNA-Seq of formalin-fixed paraffin-embedded (FFPE) tissue

**DOI:** 10.1101/2022.08.25.505257

**Authors:** Hattie Chung, Alexandre Melnikov, Cristin McCabe, Eugene Drokhlyansky, Nicholas Van Wittenberghe, Emma M. Magee, Julia Waldman, Avrum Spira, Fei Chen, Sarah Mazzilli, Orit Rozenblatt-Rosen, Aviv Regev

**Affiliations:** Klarman Cell Observatory, The Broad Institute of MIT and Harvard, Cambridge, MA, USA; Division of Computational Biomedicine, Boston University School of Medicine, Boston, MA, USA; Department of Stem Cell and Regenerative Biology, Harvard University, Cambridge, MA, USA; Department of Biology, Massachusetts Institute of Technology, Cambridge, MA, USA; Genentech, 1 DNA Way, South San Francisco, CA, USA

## Abstract

Profiling cellular heterogeneity in formalin-fixed paraffin-embedded (FFPE) tissues is key to characterizing clinical specimens for biomarkers, therapeutic targets, and drug responses. Here, we optimize methods for isolating intact nuclei and single nucleus RNA-Seq from FFPE tissues in the mouse brain, and demonstrate a pilot application to a human clinical specimen of lung adenocarcinoma. Our method opens the way to broad applications of snRNA-Seq to archival tissues, including clinical samples.

## Main Text

High resolution profiling of the molecular and cellular heterogeneity in human clinical specimens is critical for advancing human biology, precision medicine, and drug discovery. Methods that enable scalable characterization of diverse clinical specimens are critical to understanding disease mechanisms, discovering biomarkers to help stratify patients, and identifying novel therapeutic targets as well as determining the impact of drugs. Single cell genomics has been highly successful at these tasks^1^, but is currently limited to either freshly harvested human tissues or fresh-frozen samples, profiled by single cell RNA-Seq (scRNA-Seq) or single nucleus RNA-Seq (snRNA-Seq), respectively^2^. In contrast, specimens of solid tissues routinely collected for histopathology are archived via formalin-fixed paraffin-embedding (FFPE). Recent technical innovations have advanced bulk RNA-Seq for FFPE samples, demonstrating the feasibility of polyA-based expression profiling even in heavily degraded tissues^3–6^. Furthermore, while spatial transcriptomics methods have increasingly enabled molecular profiling of FFPE specimens, these methods are not at single cell resolution and have limited detection of genes^7^. Thus, scalable single-cell profiling of FFPE samples remains a challenge^8^. FFPE tissues pose numerous difficulties for applying single-cell genomics, including the extraction of intact cells or nuclei from damaged cellular structures, and detecting heavily degraded, low quantity RNA^6^. In particular, a nucleus-based method should offer a compelling option that circumvents the challenge of dissociating intact whole cells from FFPE specimens where membranes might be too damaged for efficient recovery^9–11^.

To this end, we present snFFPE-Seq, a method for snRNA-Seq of FFPE samples, by optimizing multiple stages of the process for both plate-based and droplet-based snRNA-Seq, including: (**1**) tissue deparaffinization and rehydration, (**2**) intact nucleus extraction, and (**3**) decrosslinking and deproteinization. We first developed snFFPE-Seq for mouse brain samples, and then applied it as a proof-of-concept to a clinical sample of human lung adenocarcinoma. We account for the reduced complexity of snFFPE profiles by computational integration with existing snRNA-Seq atlases from frozen specimens of the same tissues^12^. To circumvent data sparsity in a human clinical snFFPE-Seq sample, we present a new computational approach, Gene Aggregation across Pathway Signatures (GAPS), that obtains more robust signals by aggregating gene counts in individual nuclei using previously defined pathway signatures.

We first developed a protocol for extracting intact nucleus suspensions from FFPE samples of the mouse brain by optimizing the deparaffinization and rehydration of tissues, then applying an established nucleus extraction method. We worked with 50 μm scrolls of the cortex area cut on a microtome to provide ideal reaction volumes and nucleus counts. We tested three deparaffinization treatments: mineral oil with heat (80°C)^13^, xylene with heat (90°C), and xylene at room temperature^14^, each followed by tissue rehydration with graded ethanol washes (**Fig. 1a**; **Methods**). We then extracted nuclei using our previously-developed lysis buffer^2,15^ that maintains the attachment of ribosomes to the nuclear membrane, thus increasing the number of captured RNA molecules. We confirmed the successful isolation of intact nucleus suspensions with transmission electron microscopy, showing that the nuclear envelope was preserved with ribosomes attached, a condition that should improve RNA capture^15^ (**Fig. 1b**).

**Figure 1.**
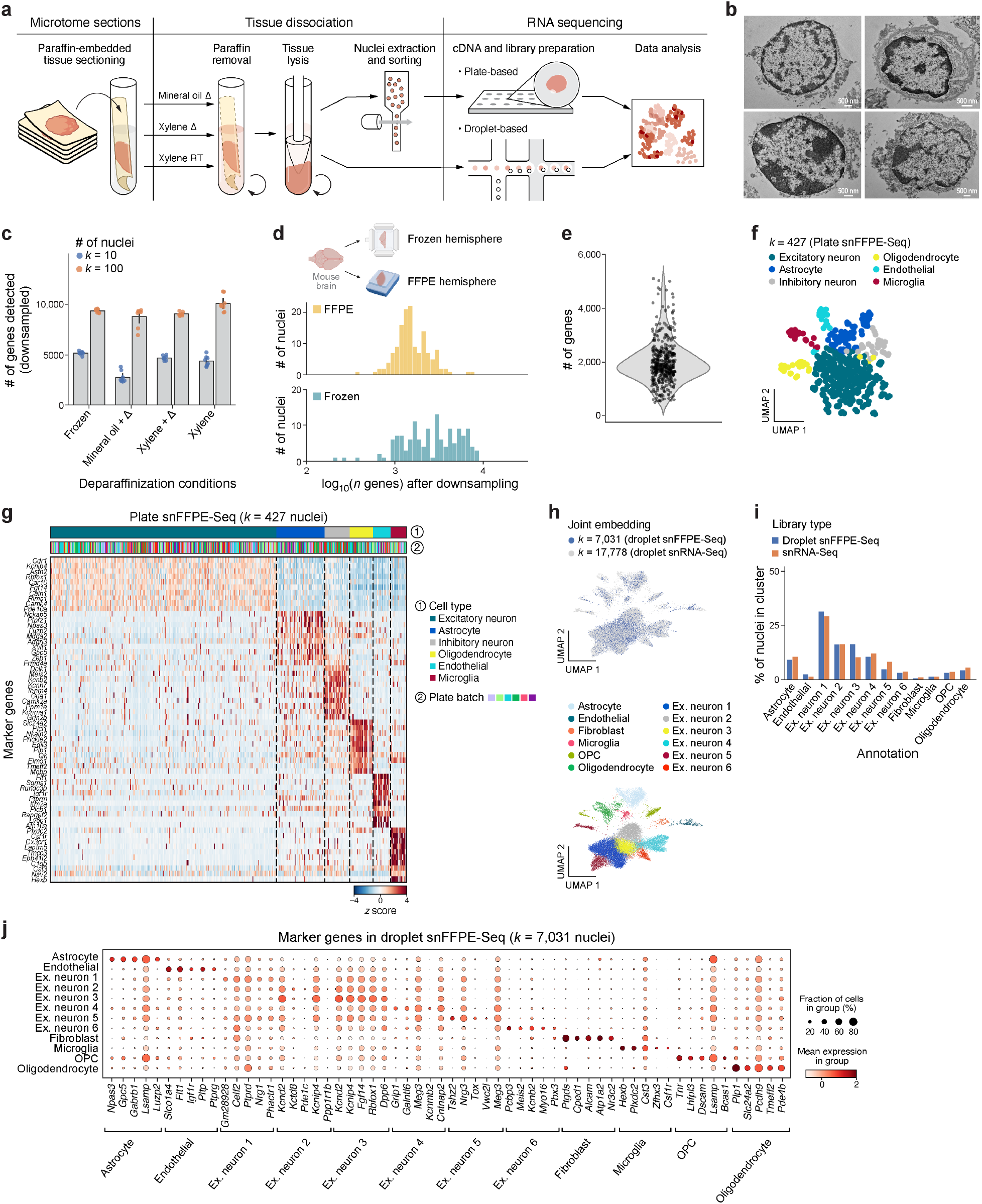
Development of single nucleus FFPE-Seq (snFFPE-Seq) in the mouse brain. **a.** Overview of the snFFPE-Seq workflow. **b.** Intact nucleus extraction from FFPE. Transmission electron microscopy of four representative intact nuclei with attached ribosomes extracted from an FFPE sample of the mouse brain, after deparaffinization with xylene at room temperature. Scale bar, 500 nm. **c.** Impact of three deparaffinization treatments on RNA capture. Number of genes (*y* axis) detected by RNA-Seq of bulk nuclei (*k*=10 or *k*=100) using SCRB-Seq by each deparaffinization condition (*x* axis), after downsampling UMI counts. Each dot indicates technical replicates (*n*=8 for each bar); error bars, 1 s.d. **d.** Good but reduced transcriptome complexity in snFFPE-Seq *vs.* snRNA-Seq of matched frozen tissue of mouse brain hemispheres. Distribution of the number of genes (log_10_, *x* axis) detected in nuclei from FFPE (top) or frozen (bottom) tissue, after downsampling reads (**Methods**). **e-g.** Plate-based snFFPE-Seq distinguishes cell types in the mouse cortex. **e.** Distribution of the number of genes detected (*y* axis) in *k*=453 nuclei profiled by SMART-Seq2 after xylene RT deparaffinization (dots). **f.** Uniform Manifold Approximation and Projection (UMAP) embedding of 453 snFFPE-Seq profiles, colored by cluster and annotated *post hoc* (color legend). **g.** Expression (*z* score, color bar) of the top 10 marker genes (rows) identified for each cluster in (f). Each nucleus profile (columns) is labeled by the corresponding cell type (top bar) and plate batch (middle bar). **h.** Cell types from the adult mouse cortex identified by joint embedding of droplet-based snFFPE-Seq and snRNA-Seq. UMAP embedding of single nucleus RNA profiles from snFFPE-Seq (*k*=7,031) and four snRNA-Seq experiments from a published study^22^ (*k*=17,778), colored by cluster and annotated *post hoc* (color legend) (left) or by profiling method (right). “Ex”: excitatory neuron. **i.** Droplet-based snFFPE-Seq and snRNA-Seq capture similar distribution of cell types. Percent of nuclei (*y* axis) from each assay (color) in each cluster (*x* axis) in (h). **j.** Marker gene expression in droplet-based snFFPE-Seq. Mean expression (log normalized counts, dot color) and proportion of expressing cells (dot size) of marker genes (columns) in each group used for annotating cell type clusters (rows) in droplet-based snFFPE-Seq nucleus profiles.

Because formalin fixation leads to extensive cross-linking of RNA to other macromolecules that pose a challenge to capturing and sequencing RNA, we reverse the cross-linking by a combination of heat^16^ and protease digestion^17^, which are compatible with plate-based RNA-Seq approaches (SMART-Seq2^18^ (SS2) and SCRB-Seq^19^). We first compared the impact of each deparaffinization treatment on bulk nuclei using SCRB-Seq, because SCRB-Seq incorporates unique molecular identifiers (UMIs) that enable assessing the efficiency of capturing unique RNA molecules. Xylene, either with heat or at room temperature, yielded a higher number of detected UMIs and genes than mineral oil (**Fig. 1c**). We chose xylene at room temperature for deparaffinization as it is easier and safer to work with than with heat. After choosing a deparaffinization condition based on UMI-based comparisons, we used SMART-Seq2^2,15^ for subsequent plate-based snFFPE-Seq experiments because SMART-Seq2 generally detected a higher number of genes than SCRB-Seq (**Supplementary Fig. 1a**).

We next assessed the difference in RNA complexity between matched FFPE treated and fresh frozen tissue from the mouse brain. Mouse cortex of both brain hemispheres of the same mouse was harvested, with one hemisphere frozen and the other treated as FFPE. From each, we extracted nuclei and profiled them individually by SMART-Seq2. FFPE nuclei profiles had ~2.7X fewer genes detected than those from the frozen sample (median genes detected: 4,382 in frozen *vs.* 1,635 in FFPE; **Supplementary Fig. 1b**), and ~2X fewer detected genes when accounting for slight variations in sequencing depth by downsampling reads (median genes: 2,927 in frozen *vs.* 1,473 in FFPE; **Fig. 1d; Methods**). The fraction of reads mapping to the reference mouse genome was lower for FFPE nuclei (median 94.1%) than for frozen nuclei (median 98.5%; **Supplementary Fig. 1b**, P=4*10^-13^, Mann-Whitney U test), as expected from degraded RNA. Mitochondrial content was <1% in both conditions (**Supplementary Fig. 1c**). Thus, while snFFPE-Seq yields fewer detected genes and mapped reads, untargeted snRNA-Seq from mouse brain FFPE still captured a substantial number of genes from the mouse transcriptome.

SnFFPE-Seq of the mouse cortex captured the expression of known cell-type marker genes. We next obtained 630 snFFPE-Seq profiles from the brains of two mice using SMART-Seq2 (**Methods**). Because FFPE samples are typically contaminated with nucleic acids from other species^20^, we aligned reads to a joint mouse (mm10) and human (hg19) pre-mRNA reference genome^21^. The majority (68%) of nucleus profiles were highly species-specific and of good quality, with >90% of reads mapped to the mouse genome (88% of nuclei), low (<5%) mitochondrial content (99% of nuclei), good (>300) gene count (84% of nuclei), and unlikely to be doublets (<450,000 counts and <5,000 detected genes; 89% of nuclei; **Supplementary Fig. 1d; Fig. 1e**). Unsupervised clustering of 427 high-quality single nucleus profiles revealed distinct subsets that reflected the expression of established marker genes (**Fig. 1f,g**; **Methods**). For example, subsets could be distinguished by marker expression as *Plp1^+^* (oligodendrocytes), *Gria1^+^Grin2b^+^* (inhibitory neurons), *Csf1r^+^Cx3cr1^+^* (microglia), and *Igf1r^+^Ly6c1^+^* (endothelial cells), among others, that were well-mixed across technical batches (**Fig. 1g**, middle bar).

Encouraged by the detection of seemingly distinct cell types, we next developed a more scalable approach that could be compatible with droplet-based platforms. Formalin can be decrosslinked by either heat (*e.g.*, during reverse transcription incubation^12^), protease digestion, or their combination. While protease-based deproteinization cannot occur inside droplets as it will degrade the reverse transcriptase enzyme, a recent study reported successful Proteinase K digestion of paraformaldehyde (PFA)-fixed *cells* before loading onto a droplet-based platform, with minimal leakage as determined by a barnyard experiment^17^. We applied a variation of this approach to FFPE nuclei by using thermolabile Proteinase K to deproteinize and decrosslink nucleus suspensions extracted from FFPE tissue at room temperature, then simultaneously heat inactivating the proteinase and partially decrosslinking the nuclei before loading onto a droplet-based platform (**Methods**). Because protease treatment reduced nucleus yield, we recommend starting with a large number (>10^5^) of nuclei, if possible.

Droplet-based snFFPE-Seq of the mouse cortex recovered broad cell types (**Fig. 1h**), although RNA damage and degradation reduced the complexity of RNA profiles as expected (**Supplementary Fig. 1e,f**). To improve cluster resolution, we co-embedded the RNA profiles from snFFPE-Seq with those from a snRNA-Seq study of the mouse cortex^22^, following a strategy we employed initially for the analysis of inCITE-Seq^12^ (a method for joint profiling of nuclear proteins and RNA in fixed nuclei, where we encountered similar challenges). We obtained robust integration of expression profiles across clusters consistent with known cell types and matching proportions across both methods (**Fig. 1i,j; Supplementary Fig. 1g**).

We next tested whether snFFPE-Seq could be applied to a clinical sample of a tumor. Using plate-based snFFPE-Seq, we collected 432 single nucleus profiles from an FFPE sample of human lung adenocarcinoma (LUAD) obtained from the primary tumor (**Fig. 2a**). For nuclei contaminated with mouse transcripts (**Supplementary Fig. 2a,b**), we removed mouse reads prior to further analysis. After filtering, we retained *k*=310 nuclei profiles for further analysis (**Supplementary Fig. 2c; Methods**). Due to the sparsity of transcriptomic profiles (**Fig. 2b**; median of 574 detected genes per nucleus), we did not perform unsupervised clustering. Instead, we classified each nucleus to a putative cell type based on known marker genes from a snRNA-Seq atlas of the healthy human lung^23^ (**Fig. 2c**; **Methods**); we prioritized using a nucleus-based atlas rather than a disease-matching cell atlas for annotation, as at the time of this writing there are no available LUAD snRNA-Seq data (only single *cell-based* data, which are challenging to integrate with *nucleus-based* data^24^). To validate cell type assignments, we reciprocally analyzed the expression of marker genes enriched in the snFFPE-Seq profiles of each assigned cell type, finding strong agreement for endothelial, epithelial, fibroblast, and muscle cells but reduced distinction between myeloid cells and lymphocytes (**Fig. 2d,e Supplementary Fig. 2d**). Expression of *EGFR, BRAF*, and *ALK*, critical targets for targeted therapy in non-small cell lung cancer^25^, was sparsely detected across assigned cell types, as expected (**Supplementary Fig. 2e**).

**Figure 2.**
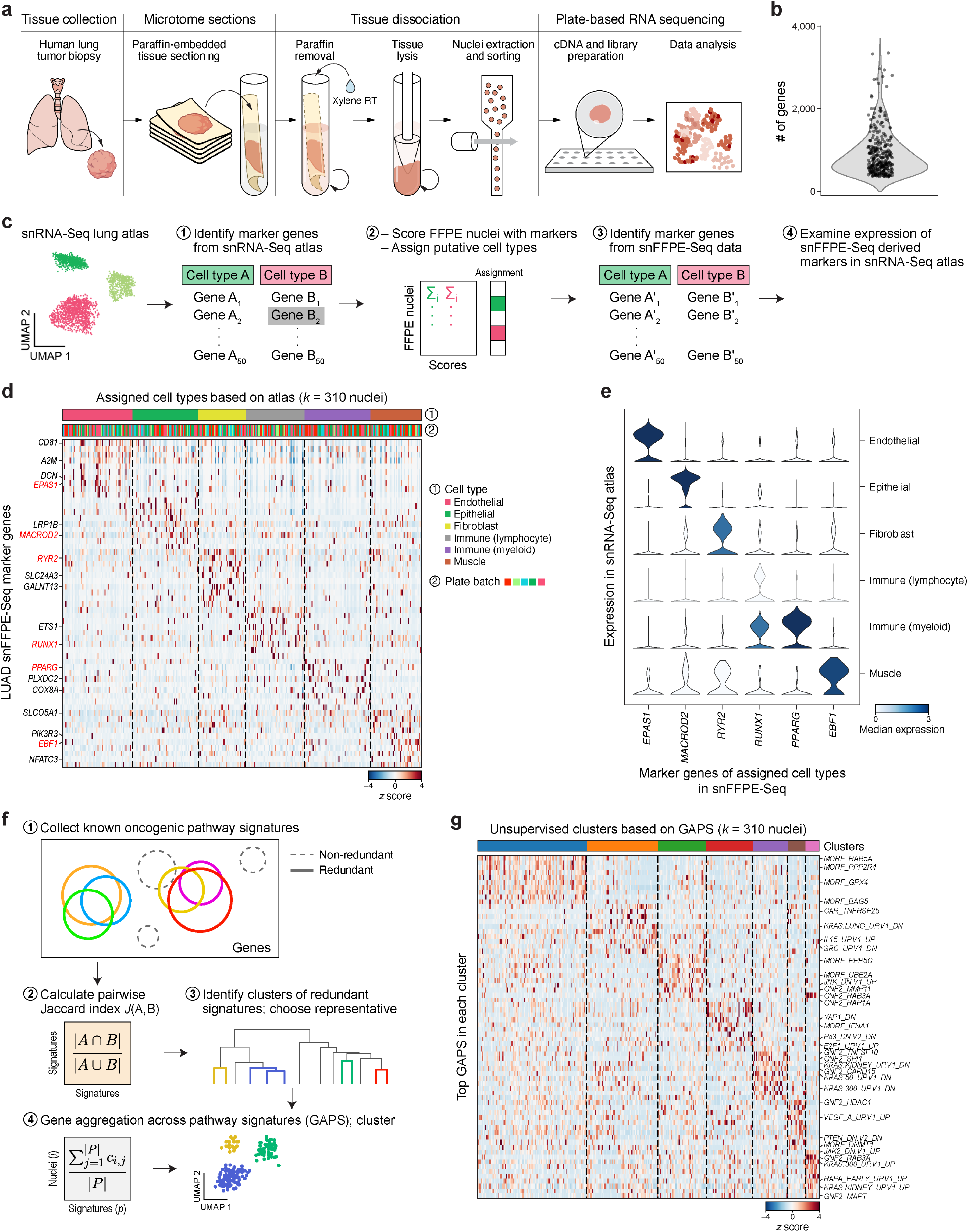
Application of snFFPE-Seq to a human lung adenocarcinoma sample. **a.** Plate-based snFFPE-Seq of a human lung adenocarcinoma (LUAD) sample. **b.** Distribution of the number of unique genes detected (*y* axis) across *k*=310 nucleus profiles (dots). **c-e.** Cell type annotation and signature detection in sparse LUAD snFFPE-Seq by atlas-based classification and annotation. **c.** Schematic of classification. **d.** Expression (color, *z* score) of top marker genes (genes, rows) corresponding to known cell types of the human lung across all profiled nuclei (columns), labeled by annotated cell type (color legend and bar) and plate batch (bottom color bar). **e.** Distribution of expression in snRNA-Seq lung atlas of select cell type marker genes (*x* axis) identified in snFFPE-Seq data for each cell type (rows) (all genes shown in **Supplementary Fig. 2d**). Color is proportional to the median expression in each set of nuclei. **f,g.** Clustering sparse snFFPE-Seq profiles by Gene Aggregation across Pathway Signatures (GAPS). **f.** Overview of strategy. **g.** Mean expression (*Z* score) of top pathway signatures (rows) in each nucleus profile (columns) labeled by clusters (bottom color bar).

To demonstrate the potential for data-driven discovery with snFFPE-Seq despite data sparsity in human samples, we clustered nucleus profiles by the expression of known cancer pathway signatures from MSigDB^26^, which identified clusters with distinct tumor-related programs. To this end, we developed a computational approach called Gene Aggregation across Pathway Signatures (GAPS), where we construct an expression matrix of nuclei-by-signatures (*i.e.*, aggregated expression across signature genes). To avoid scoring the same gene sets repeatedly, we sought to identify non-redundant signatures: we clustered signatures by their pairwise gene membership Jaccard similarity scores, then selected a representative signature from each signature set (**Fig. 2f; Supplementary Fig. 3a**; **Methods**). Finally, we clustered nuclei by their per-nucleus aggregated signature profiles, identifying 7 distinct nucleus clusters. Unsupervised clusters revealed several associated with tumor-related signatures (**Fig. 2g**): one enriched for the upregulation of lung adenocarcinoma-related signature KRAS and the mTOR pathway (*e.g.*, KRAS.300_UP.V1_UP, RAPA_EARLY_UP.V1_UP), a separate cluster enriched for the downregulation of KRAS signature (*e.g.* KRAS.50_UP.V1_DN), and one reflecting signatures of the tumor suppressor PTEN and HDAC1 (*e.g.* PTEN_DN.V2_DN, GNF2_HDAC1). Thus, snFFPE-Seq can detect higher-resolution variations in tumor cell subsets.

In conclusion, snFFPE-Seq opens the way to scalable snRNA-Seq of FFPE samples, an essential sample source for clinical research. Our work provides a critical advance to profiling the vast resource of FFPE specimens, enabling greater access to the molecular diversity of human clinical samples across heterogeneous patient populations. Notably, a significant limitation to scaling is the high variability in the preparation of FFPE samples, including different formalin incubation durations and storage conditions which impact RNA quality. Furthermore, for large tissue specimens, some cells in the middle of the tissue can remain alive during fixation as formalin slowly penetrates, providing sufficient time for gene expression changes and cell death^27^. To mitigate this, we recommend quantifying the quality of bulk RNA extracted from a portion of the FFPE block before proceeding with snFFPE-Seq. For FFPE samples with heavily degraded, short RNA fragments, random primers^20^ or polyadenylation of short RNA sequences with SMART-Seq-total^28^ may improve the capture rate. Furthermore, our nucleus extraction method can be coupled to multiple other profiling methods, including multiplexed antibody-based detection of proteins^12^ or targeted mutation profiling^29,30^. Further optimization of tissue-specific snFFPE-Seq protocols combined with emerging spatial transcriptomics techniques for FFPE^8,31,32^ and new computational methods that tackle sparsity should significantly enhance our understanding of the functional organization and interactions of cells in tissues, especially in disease.

## Methods

### Human subjects

Adult patients included in this work provided preoperative informed consent to participate in the study according to Institutional Review Board protocol at Boston Medical Center H-27014.

### Mice

C57BL/6J (Jax 000664) mice were purchased from The Jackson Laboratory and bred in-house. Male and female mice were used at 8-12 weeks of age. All mice were maintained under SPF conditions on a 12-h light-dark cycle and provided food and water ad libitum. All mouse experiments were approved by and performed per the Institutional Animal Care and Use Committee guidelines at the Broad Institute.

### FFPE preparation of mouse brain

Adult 8-12 week-old mice were euthanized by CO_2_. The entire mouse brain tissue was dissected, placed in embedding cassettes, and fixed in 4% methanol-free paraformaldehyde at 4°C overnight. Fixed tissue was then dehydrated in 80% ethanol and processed on the Vacuum Infiltrating Tissue Processor (VIP) at the Koch Institute Histology Core as follows: 70% ethanol for 45 min, 85% ethanol for 45 min, 95% ethanol for 3x 45 min, 100% ethanol 2x 45 min, xylene 3x 45 min. Tissues were embedded into paraffin wax at 58-60°C across four changes, 30 min each. For histology, FFPE blocks were sectioned at 20 μm and stained with hematoxylin and eosin. All FFPE tissue samples were prepared weeks before testing; older blocks, especially those not optimally preserved, are more likely to have degraded RNA.

### FFPE preparation of human lung adenocarcinoma

FFPE lung tissue samples were obtained from Boston Medical Center (BUMC). Briefly, a resection of human lung adenocarcinoma was processed with standard histopathology procedure for 24 hours in 10% neutral buffered formalin (NBF) before processing through graded ethanol dehydration steps, embedded in paraffin, then stored at room temperature. FFPE blocks were sectioned into 50 μm scrolls, collected into 1.7 mL Eppendorf tubes, and maintained at 4°C before processing for snFFPE-Seq. Starting with thinner scrolls, *e.g.*, 25 μm, resulted in the loss of a pellet during nucleus extraction.

### Deparaffinization, tissue rehydration, and nucleus extraction

FFPE blocks were prepared by cutting 50 μm scrolls on a microtome (cleaned with 70% EtOH and RNAseZAP) and stored in a sterile safe lock 1.5 mL Eppendorf tube at 4°C. Deparaffinization of 50μm scrolls was tested with three different methods. Each protocol started with three 50 μm FFPE scrolls placed in a 1.5 mL safe lock Eppendorf tube. Excess paraffin was trimmed with a razor blade.

1. *Mineral oil with heat.* We added 500 μl of mineral oil to the tube with FFPE scrolls and incubated at 80°C for 5 min on a heat block. After a quick vortex and spin in a microcentrifuge, 750 μl of 95% ethanol was added and incubated for 2 min at 80°C. The tube was spun down to create a phase separation, and the upper phase of the mineral oil was removed thoroughly. The tissue was resuspended with 1 mL of 95% ethanol pre-warmed at 80°C, vortexed, and incubated at room temperature (RT) for 2 min. After a spin down, residual oil drops in the upper phase and the ethanol were removed. We then conducted the following ethanol rehydration steps at RT: twice with 1 mL of 75% ethanol, and twice with 1 mL of 50% ethanol.
2. *Xylene with heat.* We added 1 mL of xylene to the tube with FFPE scrolls, incubated at RT for 10 min, and spun down. Xylene was removed, and the process was repeated twice but with 10 min incubations at 90°C, for a total of 3 xylene washes. Tissue was rehydrated at RT with 1 mL of 95%, 75%, and 50% ethanol with 2 min incubations each, repeating each ethanol concentration twice. The tube was spun down after each incubation and ethanol removed.
3. *Xylene at room temperature*. We added 1 mL of xylene to the tube with FFPE scrolls, incubated at RT for 10 min, and spun down in a microcentrifuge. Xylene was removed, and the process was repeated twice for a total of 3 xylene washes. Tissue was rehydrated at RT with 1 mL of 95%, 75%, and 50% ethanol with 2 min incubations each, repeating each ethanol concentration twice. The tube was spun down after each incubation and ethanol removed.

Nucleus extraction was performed after deparaffinization and rehydration with the following protocol as previously developed^2,15^. All nucleus extractions were conducted on ice and/or at 4°C.

2X salt-Tris (ST) buffer: 292 mM NaCl (ThermoFisher #AM9760G), 20 mM Tris-HCl pH 7.5 (ThermoFisher #15567027), 2 mM CaCl_2_ (Sigma Aldrich #21115), 42 mM MgCl_2_ (ThermoFisher #AM9530G) in ultrapure water (ThermoFisher #10977015).
1X Salt-Tris buffer without MgCl_2_ (ST-): 146 mM NaCl, 10 mM Tris-HCl pH 7.5, 1 mM CaCl_2_ in ultrapure water, and 40 U/mL SUPERaseIn (ThermoFisher #AM2696).
1X ST buffer: 1 part 2X ST buffer, 1 part ultrapure water, 40 U/mL SUPERaseIn.
CST lysis buffer (scaled appropriately): 1 mL of 2X ST buffer, 980 μl of 1% CHAPS (Millipore #220201), 10 μl of 2% BSA (NEB B9000S), 2 μl of 20 U/mL SUPERaseIn, 8 μl ultrapure water.

#### Mouse brain

On ice, rehydrated tissue was placed into a glass douncer (Sigma Aldrich D8938) with 2 mL of ice-cold CST lysis buffer, then dounced 25x with pestle A followed by 25x with pestle B. The homogenized lysate was passed through a 30 μm filter (Miltenyi #130-041-407). An additional 2 mL of 1X ST buffer was used to rinse the douncer, then passed through the filter. A final 1 mL of 1X ST buffer was added to bring the final volume to 5 mL, and incubated on ice for 5 min. The tube was spun at 500g for 5 min at 4°C in a swinging bucket centrifuge. After removing the supernatant, the pellet was resuspended in 1 mL of 1X ST buffer, incubated on ice for 5 min, spun at 500g for 5 min at 4°C, and resuspended in 500 μl of 1X ST buffer. An aliquot of nuclei was stained with DAPI and counted under a fluorescent microscope.

#### Human lung adenocarcinoma

On ice, rehydrated tissue was placed into a well of a 6-well plate (Stem Cell Technologies #38015) with 1 mL of ice-cold CST lysis buffer. The tissue was finely chopped using Noyes Spring Scissors (Fine Science Tools #15514-12) for 10 min on ice. The homogenized lysate was filtered through a 40 μm Falcon cell strainer (ThermoFisher #08-771-1) into a 50 mL Falcon tube. Another 1 mL of cold CST was used to wash the well and added through the filter. The volume was brought up to 5 mL with 3 mL of 1X ST buffer, transferred to a 15 mL Falcon tube, and incubated on ice for 5 min. Nuclei extract was spun at 500g for 5 min at 4°C in a swinging bucket centrifuge. After removing the supernatant, the pellet was resuspended in 500 μl 1X ST buffer and filtered through a 35 μm Falcon cell strainer (Corning #352235). An aliquot of nuclei was stained with DAPI and counted under a fluorescent microscope.

#### Fluorescence-activated cell sorting (FACS) for plate-based sequencing

Nucleus suspensions were stained with Vybrant DyeCycle Ruby (ThermoFisher #V10309) at 1:500 and filtered through a 20 μm filter (Miltenyi #130-101-812). Individual nuclei were sorted on a Sony Sorter SH800 with a 100 μm sorting chip into wells of a 96-well plate containing 5 μl of Buffer TCL (Qiagen #1031576) with 1% β-mercaptoethanol (ThermoFisher #21985023).

### Single nucleus RNA-Sequencing with deproteinization and decrosslinking

#### Plate-based SCRB-Seq and SMART-Seq2

Plate-based snFFPE-Seq protocols were carried out by adding 1 μl of 1 μg/μl proteinase K (ThermoFisher #AM2548) to each well with the sorted FFPE nuclei, followed by incubation at 55°C for 15 min, then crosslink reversal at 80°C for 15 min. Post incubation cleanup was conducted using 2.2X by volume of Agencourt RNAClean XP beads (Beckman Coulter, #A63987) used according to the manufacturer’s protocol. All subsequent steps, including library construction, were carried out following the standard SCRB-Seq^33^ and SMART-Seq 2^34^ protocols, except reverse transcription reactions were enhanced by increasing MgCl_2_ concentration to 10 mM and by the addition of trehalose (Life Sciences #TSIM100) to 0.6 M.

SCRB-Seq libraries were sequenced on a NextSeq 500/550 with 16 cycles for read 1, 8 cycles for index 1, and 68 cycles for read 2. SMART-Seq 2 libraries were sequenced on a NextSeq 500/550 with 38 cycles for read 1, 8 cycles for index 1, 8 cycles for index 2, and 38 cycles for read 2.

#### Droplet-based scRNA-Seq

Nucleus suspensions were adjusted to ~10^4^ nuclei/μl in 100 μl of 1X ST(-) buffer. To deproteinize, 2 μl of undiluted Thermolabile Proteinase K (NEB #P8111S) and 1 μl SUPERaseIN (20 U/μl) were added to the suspension and incubated for 30 min at room temperature, followed by proteinase inactivation and reverse crosslinking for 10 min at 55°C on a heat block. Nuclei extract was spun at 500g for 5 min at 4°C in a swinging bucket centrifuge. After removing the supernatant, the pellet was resuspended in 100 μl of ice-cold 1X ST(-) buffer. The nuclei were then placed on ice, counted, and adjusted appropriately to a concentration of ~10^3^ nuclei/ μl for loading the 10X Chromium chip. We loaded 15,000 nuclei onto a single channel of the Chromium Chips for the Chromium Single Cell 3’ Library (V3, PN-1000075). All subsequent steps, including library construction, were prepared according to the standard protocol according to the manufacturer’s instructions. Libraries were sequenced on a HiSeq X with 28 cycles for read 1, 8 cycles for index 1, and 96 cycles for read 2.

### Data pre-processing

Plate-based data were pre-processed with the zUMIs pipeline^35^ version 2.4.5b (for SMART-Seq2 and SCRB-Seq), and droplet-based data were pre-processed with CellRanger version 3.1.0 on Cumulus version 1.0^36^. Reads from demultiplexed FASTQ files were aligned to pre-mRNA annotated genomes of the jointly combined mouse (mm10) and human (hg19) reference genomes as previously described^21^. All reads were aligned to the mm10_and_hg19_premRNA reference genome^21^.

### Comparison of gene counts across deparaffinization protocols

To compare RNA capture across deparaffinization, UMIs were pooled from all nuclei profiles in one snFFPE-Seq or snRNA-Seq experiment, and down-sampled to the minimum number of UMIs detected in frozen nuclei: 18,159 UMIs for *k*=10 nuclei and 59,608 UMIs for *k*=100 nuclei.

### Comparison of gene counts between snFFPE-Seq and snRNA-Seq

To compare the number of genes between snFFPE-Seq and snRNA-Seq of mouse brain using SMART-Seq2, reads were downsampled to the median counts detected among FFPE nuclei (47,887 counts).

### Clustering of SMART-Seq2 snFFPE-Seq of mouse brain

All analyses were conducted with scanpy v1.9.1^37^. Nucleus profiles were retained if and only if >90% of their detected genes were mapped to the mouse (mm10) reference, <5% of reads were mitochondrial, at least 300 detected genes, and no more than 450,000 counts and 5,000 genes. Raw counts were normalized by ln(gene length), then normalized per nucleus using scanpy’s normalize_per_cell function, and ln+1 transformed. Of the 20,347 genes detected, 2,059 highly variable genes were selected using the highly_variable_genes function in scanpy (min_mean=0.32, max_mean=2, min_disp=0.5). The number of mouse genes detected was regressed, followed by plate batch, and data were clipped at max_value=10. Dimensionality reduction was performed using Principal Component Analysis (PCA), a *k-*nearest neighbor (*k*-NN) graph was constructed with the top 30 PCs and *k=10* neighbors, clustered with the Leiden algorithm^38^, and projected into a uniform manifold approximation and projection (UMAP) embedding^39^. Marker genes were identified for each cluster by comparing the nuclei profile in that cluster to profiles for all other clusters using a t-test (**Supplementary Table 1**).

### Analysis of gene expression for droplet-based snFFPE-Seq of the mouse brain

Nucleus profiles were retained if and only if <5% of reads were mitochondrial and had at least 220 but no more than 1000 detected genes. Genes detected in >2 filtered nuclei were kept. Raw counts were normalized per nucleus using scanpy’s normalize_total function and ln+1 transformed. SnFFPE-Seq nuclei profiles from this study (*k*=7,078) were jointly embedded with snRNA-Seq data of the cortex from a published study^22^ (*k*=17,948; WT only). Of the 19,905 genes detected, 5,555 highly variable genes were selected using the highly_variable_genes function in scanpy (min_mean=0.0016, max_mean=0.16, min_disp=0.31). The number of counts and the fraction of mitochondrial reads were regressed, followed by sequencing assay type (snRNA-Seq vs. snFFPE-Seq), then scaled and clipped at max_value=10. Further integration across sequencing assay types was conducted via an implementation of Harmony^40^. Dimensionality reduction was performed using Principal Component Analysis (PCA), a *k*-nearest neighbor (*k*-NN) graph was constructed with the top 40 PCs and *k*=10 neighbors, clustered with the Leiden algorithm^38^, and projected into a uniform manifold approximation and projection (UMAP) embedding^39^. A cluster with high mitochondrial content (*k*=11 nuclei) and a cluster whose top marker genes were lncRNAs and mitochondrial genes without an obvious match to known cell types of the cortex (*k*=206) were removed. The final embedding consisted of *k*=7,031 snFFPE-Seq and *k*=17,778 snRNA-Seq nuclei RNA profiles. Marker genes were identified for each cluster by comparing the nuclei profile in that cluster to profiles for all other clusters using a t-test (**Supplementary Table 2**).

### Cell type annotation of snFFPE-Seq of human lung adenocarcinoma

#### Pre-processing

All analyses were conducted with scanpy v1.9.1^37^. Nucleus profiles were retained if and only if <10% of reads were mitochondrial and had at least 200 but no more than 4,000 detected genes, yielding 310 nucleus profiles (of 432) with 16,920 human genes detected in at least one profile. Given the high expression of lncRNA, genes starting with RP11 and LINC were removed (3,783 such genes removed). The final count matrix for downstream analysis consisted of 310 nuclei and 16,556 genes. Counts were normalized within each nucleus, transformed to ln+1 counts, regressed out for the number of hg19 genes detected followed by the plate batch, then scaled with max_value at 10.

#### Assigning cell types

The top 50 marker genes of broad cell types from a previously annotated snRNA-Seq atlas of the healthy human lung (Cell types level 2)^23^ were used to calculate a cell type score for each snFFPE-Seq profile, using the score_genes function in scanpy (t-test). Only marker genes detected in snFFPE-Seq were used (46 epithelial, 45 endothelial, 49 fibroblast, 47 lymphocyte, 49 myeloid, 46 muscle; **Supplementary Table 3**), and these genes were used to calculate a cell type score for each FFPE nucleus profile using the score_genes function in scanpy. Each nucleus profile was assigned a putative cell type identity based on the maximum score. We then identified genes enriched in the snFFPE-Seq data based on their assigned cell types using the rank_genes_groups function (t-test; **Supplementary Table 4**), then reciprocally examined their expression in the snRNA-Seq lung atlas^23^.

### Clustering of snFFPE-Seq by Gene Aggregation across Pathway Signatures (GAPS)

For each of 616 MSigDB tumor-related pathway signatures (c4.cgn, cancer gene neighborhoods; c6, oncogenic signature set), a signature was retained if it was comprised of at least 20 and at most 150 detected genes in the FFPE data and if at least 50% of its member genes were detected, resulting in 499 signatures (**Supplementary Table 5**). To create a signature expression *x_i,p_* for each nucleus *i*, raw counts *c_i,j_* were aggregated across the signature genes and normalized by the number of genes in the signature (that are also expressed in the dataset), |*P*|:

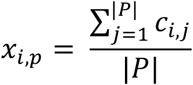

Redundant signatures were removed by the following procedure. First, the Jaccard index was calculated for each pair of signatures *A* and *B* as *J*(*A,B*) = |*A*∩*B*| / |*A*∪*B*| based on the gene sets defining each signature. Signatures were clustered by their Jaccard similarity profiles with a Euclidean distance and the ward method. The linkage matrix was used to cut the dendrogram at a threshold of 2.2, identifying 22 signature sets (**Supplementary Fig. 3a**). Sixteen of the 22 signature sets contained redundant signatures, defined if a signature set’s median within-cluster pairwise Jaccard index greater was than 0.15 (color block in **Supplementary Fig. 3a**). A representative signature was selected for each of the 16 signature sets (to preserve interpretability and annotation) as the signature with the maximum median pairwise Jaccard index within each set. The remaining 184 signatures *x_i,p_* were removed from the nuclei *x* signatures expression matrix, resulting in 308 unique GAPS (292 GAPS not in a redundant signature set and 16 representative GAPS of each of the 16 redundant sets) across 8,370 unique genes (**Supplementary Table 6**).

The filtered GAPS expression matrix was normalized and transformed to ln+1 counts. Of 308 unique GAPS, 75 highly variable GAPS were selected using the highly_variable_genes function in scanpy (min_mean=0.5, max_mean=2, min_disp=0.25, batch_key=’plate’). The number of counts (across all genes), the number of GAPS per nucleus, and the plate batch were all regressed, and then data were scaled with max_value at 10. Dimensionality reduction was performed using Principal Component Analysis (PCA), a *k-*nearest neighbor (*k*-NN) graph (*k*=10 nearest neighbors) was constructed with the top 40 PCs and *k*=10 neighbors, clustered with the Leiden algorithm, and projected into a uniform manifold approximation and projection (UMAP) embedding. Marker GAPS were identified for each cluster by comparing the nucleus profiles in that cluster to profiles for all other clusters using a t-test (**Supplementary Table 7**).

## Data Availability Statement

Gene expression count matrices and raw FASTQ files for all **mouse** snFFPE-Seq data have been uploaded to Gene Expression Omnibus under accession GSE211797. Gene expression counts and raw FASTQ files of the human lung adenocarcinoma sample will be uploaded on a controlled access platform. Mouse cortex data^22^ used for the joint embedding of the mouse cortex is available under GSE143758. Human snRNA-Seq atlas data^23^ used to annotate the lung data is available at https://gtexportal.org/home/datasets.

## Code Availability

All code used for analyses is available at https://github.com/klarman-cell-observatory/snFFPE-Seq.

## Acknowledgments

We thank Dr. Eric Burks at Boston Medical Center for providing the LUAD clinical samples. We thank the Koch Institute Histology Core, the Harvard Medical School Electron Microscopy Facility, and the Broad Institute Flow Cytometry Core. We thank L. Gaffney and A. Hupalowska for assistance with figures. This research was supported in part by the Klarman Cell Observatory, the National Cancer Institute, and by the Human Tumor Atlas Pilot Project (HTAPP). A.S. and S.M. were supported by NCI U01CA196408. The funders had no role in study design, data collection and analysis, decision to publish, or preparation of the manuscript.

## Author Contributions

H.C., A.M., C.M., E.D., O.R-R., A.R. designed the study. H.C., A.M., C.M., N.V.W., E.M.M., J.A. conducted experiments with supervision from O.R-R. and A.R. H.C. harvested mouse brain samples. A.S. and S.M. provided human lung adenocarcinoma samples. H.C. conducted analyses with supervision from A.R. H.C. and A.R. wrote the paper with input from all authors.

---

**Supplementary Figure 1.**
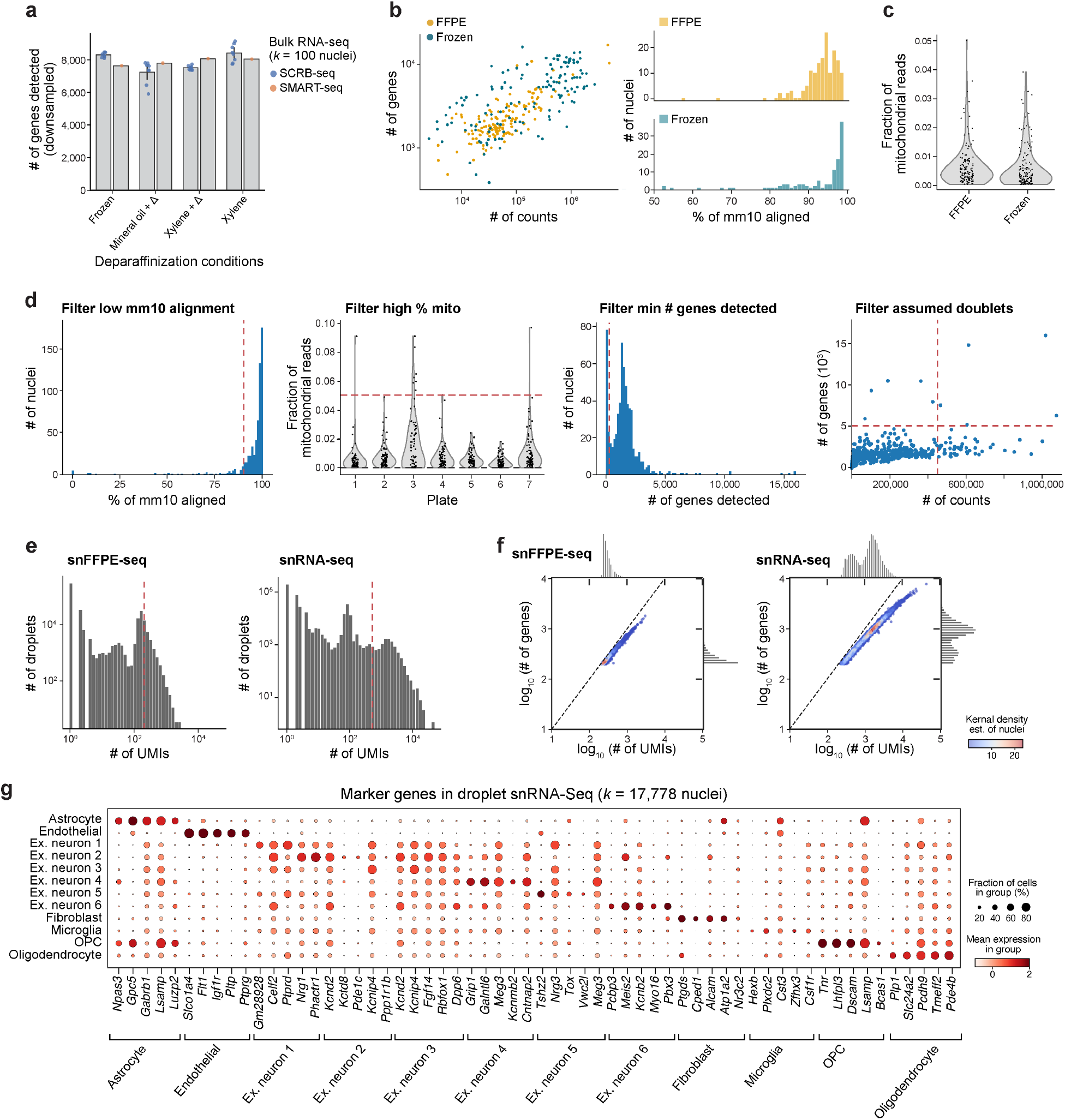
Quality control metrics related to the development of snFFPE-Seq in the mouse brain. **a.** Comparing SCRB-Seq *vs.* SMART-Seq2-Seq for RNA detection across deparaffinization conditions. Number of genes detected (*y* axis) in bulk RNA-Seq of *k*=100 nuclei extracted from frozen or FFPE mouse brain across deparaffinization conditions (*x* axis) with SCRB-Seq or SMART-Seq2-Seq (color). Reads were downsampled to 50,000 per sample to conduct a fair comparison. Each dot indicates technical replicates (*n*=8 for SCRB-Seq, *n*=1 for SS2), where each replicate is across *k*=100 nuclei. Error bars, 1 s.d. **b,c.** Comparison of RNA profile quality metrics in frozen vs. FFPE nuclei from matching hemispheres of the same mouse brain. **b**. *Left:* Number (log_10_) of unique genes (*y* axis) and reads (*x* axis) detected in individual nuclei (dots) colored by tissue treatment. *Right*: Distribution of the fraction (%) of reads aligned to the mm10 genome (*x* axis) by tissue treatment (FFPE, top; frozen, bottom). **c.** Distribution of the fraction of mitochondrial reads (*y* axis) in each nuclei profiled from FFPE or frozen tissue (*x* axis). **d.** Quality control metrics and thresholds used to select high quality nuclei profiles from plate-based snFFPE-Seq. From left to right: Distributions of fraction (%) of reads aligned to the mouse genome in each nucleus (*x* axis, left) or to the mitochondrial genome (*y* axis) in each plate (*x* axis) (second from left); of the number of genes (*x* axis, second from right) detected in each nucleus; and the number of counts (*x* axis) and the number of genes (*y* axis) in each nucleus to filter suspected doublets (far right). Red lines indicate thresholds used to filter nuclei and the label on top indicates the direction of the filter. **e-g.** Quality measures for droplet-based snFFPE-Seq of the mouse cortex. **e.** Distribution of number of UMIs (x axis) in each droplet from snFFPE-Seq (left) or published snRNA-Seq (right). Red line: threshold used to filter. f. Distributions (marginals) of the number of UMIs (*x* axis) and genes (*y* axis) from snFFPE-Seq (left) and published snRNA-Seq (right). Density of individual nuclei (dots) is calculated with a Gaussian kernel estimate. **g.** Mean expression (log normalized counts, dot color) and fraction of expressing cells (dot size) of select marker genes (columns) in nuclei of each cell type (rows) in snRNA-Seq (top).

**Supplementary Figure 2.**
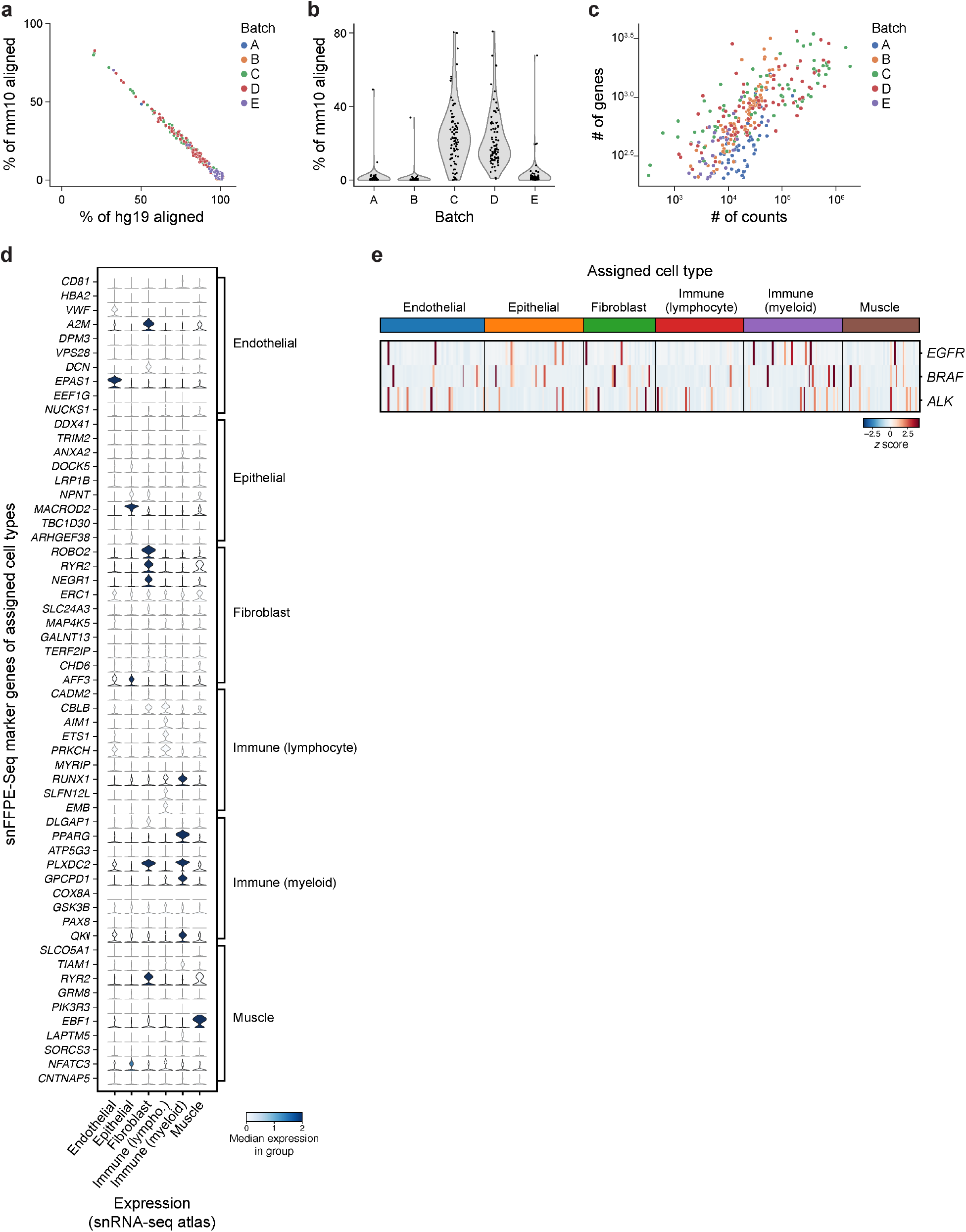
Quality control metrics and characterization of snFFPE-Seq of a human lung adenocarcinoma (LUAD) sample. **a-c.** Quality characteristics across batches. **a.** Percentage of reads aligned to the mouse (mm10, *y* axis) and human (hg19, *x* axis) genomes, in each nucleus (dots) colored by plate batch (color). **b.** Distribution of % reads aligned to the mouse genome (*y* axis) for individual nuclei (dots) in each plate batch (*x* axis). **c.** Number of unique genes detected (log_10_, *y* axis) and number of unique reads (log_10_, *x* axis) for each nucleus (dots) colored by plate batch. **d.** Marker gene expression of assigned cell types derived from snFFPE-Seq, shown in snRNA-Seq atlas of the healthy human lung. Distribution of expression in snRNA-Seq lung atlas of each cell type marker gene (*x* axis) identified in snFFPE-Seq data for each cell type (rows). Color is proportional to the median expression in group. **e.** Lung cancer driver oncogene expression in snFFPE of LUAD. Expression (colorbar, *Z* score) of *EGFR, BRAF, ALK* (rows) across LUAD snFFPE-Seq nucleus profiles (columns), grouped by assigned cell type (color).

**Supplementary Figure 3.**
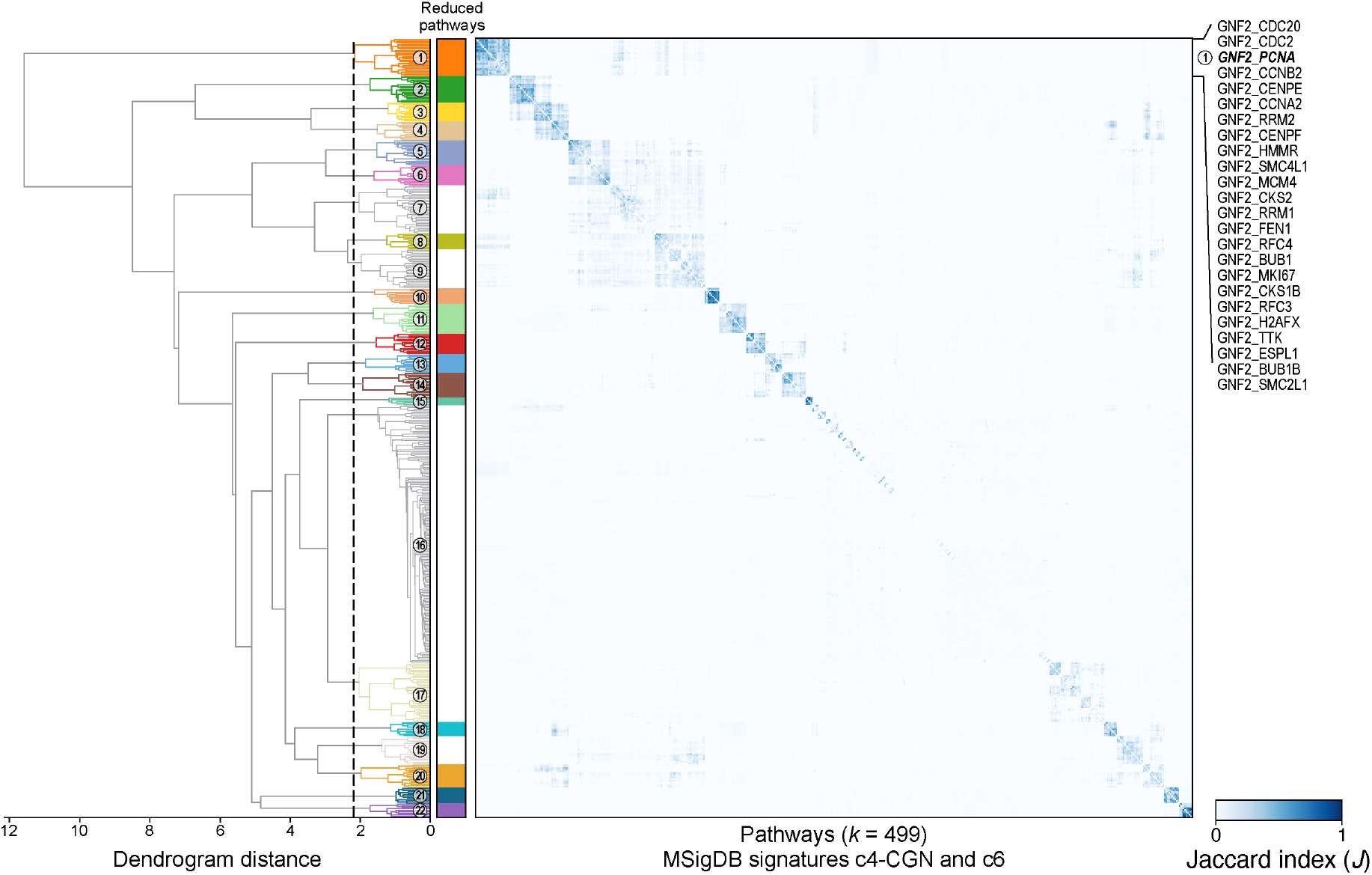
Identifying redundant pathway signatures. Jaccard similarity index (color) of each pair of pathway signatures (rows, columns), hierarchically clustered with a Euclidean distance and the ward metric. Dashed line: dendrogram (left) cut at a distance threshold of 2.2. Colored numbered branches: leaf assignment to 22 clusters based on their cut branch. Matching color block: Clusters of signatures considered as redundant (median pairwise Jaccard index > 0.15).

